# Microplastic vector effects: are fish at risk when exposed via the trophic chain?

**DOI:** 10.1101/2019.12.17.879478

**Authors:** Agathe Bour, Joachim Sturve, Johan Höjesjö, Bethanie Carney Almroth

**Affiliations:** Department of Biological and Environmental Sciences, University of Gothenburg, Göteborg, Sweden

**Keywords:** ecotoxicity, chlorpyrifos, behavior, uptake, stickleback

## Abstract

In aquatic organisms, trophic transfer is a relevant exposure route for microplastics (MPs). Despite their relevance, effect studies on fish exposed via trophic chains are currently very scarce. MPs are known to contain many chemicals that could be transferred to organisms and induce deleterious effects. However, there is currently no consensus on whether MPs represent a significant exposure pathway to chemicals in contaminated habitats. Here, we exposed three-spined sticklebacks (*Gasterosteus aculeatus*) to polyethylene MPs via prey ingestion, in a one-month experiment. MPs were either pristine or spiked with chlorpyrifos (CPF), and a CPF control was included to compare vector effects of MPs and natural prey. Following exposure, we assessed AChE activity and fish behavior (feeding, locomotion, environment exploration and reaction to the introduction of a novel object). No effect was observed in fish exposed to pristine MPs. CPF accumulation was observed in fish exposed to CPF-spiked MPs (MP-CPF), confirming the vector potential of MPs. However, CPF accumulation was more important in fish exposed to CPF via prey. In fish exposed to MP-CPF, we observed significant AChE inhibition and hyperactivity, which could result in increased vulnerability to predation. CPF organ distribution differed between groups, suggesting that chemical exposure via MPs could alter organ distribution of chemicals. This can result in a change in the organs most at risk, likely increasing intestine exposure.

## 1. Introduction

Increasing numbers of field and laboratory studies have shown that most lower trophic level organisms are able to ingest microplastics (MPs) (Lusher, 2015; Scherer et al., 2018). Ingestion of MP-contaminated prey by predator species is therefore very likely and trophic transfer has been identified as a relevant contamination pathway for MPs (Farrell and Nelson, 2013; Nelms et al., 2018). Despite their relevance, MP trophic transfer and its impacts on upper trophic level organisms are still poorly investigated, especially in studies involving fish. MPs are known to contain many additives, such as plasticizers, flame retardants, stabilizers, surfactants and pigments (Lambert and Wagner, 2018), as well as environmental contaminants, such as PCB, PAHs and PBDEs (Koelmans, 2015). A major concern is therefore their potential to act as vectors and to transfer chemicals to organisms (Rochman, 2019). However, modelling studies have questioned this hypothesis, arguing that the role of MPs in chemical transfer to organisms could be minor in a context of contaminated environments (Teuten et al., 2007; Gouin et al., 2011; Koelmans et al., 2013). Especially, the ingestion of contaminated prey and/or natural particles could result in greater chemical uptake, compared to MPs. Despite the importance of comparing MPs to other natural vectors of contamination, such as natural prey, these alternative exposure pathways are still poorly investigated in MP ecotoxicity studies (Koelmans, 2015), especially those focusing on aquatic organisms.

To explore these major knowledge gaps, the present study aims to investigate i) the effects of MPs on fish exposed via prey ingestion, ii) the potential of MPs to transfer chemicals (*i.e.* vector effect) to fish exposed via prey ingestion, iii) the relative importance of MPs’ vector effect, in comparison with contaminated prey, and iv) the consequences of MPs’ vector effect on organisms’ performance. For this purpose, we studied an experimental trophic chain comprising the three-spined stickleback (*Gasterosteus aculeatus*) as predator species, and brine shrimps (*Artemia sp.*) as natural prey. The three-spined stickleback naturally occupies a wide range of aquatic habitats (Bell and Foster, 1994) and has been widely used in ecotoxicity and behavior studies (Girvan and Braithwaite, 1998; Sturm et al., 2000; Dingemanse et al., 2007; Jutfelt et al., 2013; Fürtbauer et al., 2015; Thompson et al., 2016; Marchand et al., 2017). For this study, stickleback individuals were fed during four weeks with brine shrimps previously exposed to pristine MPs, chemical-spiked MPs or chemical-contaminated water. To study the vector effect of MPs, we selected polyethylene (PE) as a model plastic polymer. PE is the most produced polymer type (PlasticsEurope, 2017) and is commonly found in environmental matrices (Bour et al., 2018). Chlorpyrifos (CPF) was selected as a model chemical compound. Its intermediate partition coefficient (log K_ow_ = 4.66) suggests the possibility to bind to MPs in aqueous environments, while potentially allowing for desorption in biological matrices. CPF is a commonly used organophosphate pesticide that inhibits acetylcholine esterase (AChE) (Ware, 1999), an enzyme involved in neurotransmission, which can result in behavioral disorder in fish, as previously shown in *Gambusia* (Rao et al., 2005). In addition to AChE inhibition, we assessed behavioral changes to study the effects of MPs and CPF on stickleback. Behavior is a sensitive endpoint likely to be affected at low contaminant concentrations, and behavior alterations can have consequences at the ecosystem level (Galloway et al., 2017). The study of behavioral endpoints is therefore very relevant in ecotoxicity studies. Here, we specially focused on feeding, locomotion and fish reaction to the introduction of a novel object.

## 2. Materials and Methods

### 2.1 Materials

#### 2.1.1 Particles and chemicals

Microplastics were purchased from Cospheric (Santa Barbara, USA; lot #120328-2--1). Opaque blue polyethylene microspheres were selected to ease quantification of exposure. According to the manufacturer, MPs are spherical (> 90% of particles), with a 27-32 μm diameter (> 90% of particles in the indicated size range) and a density of 1.00 g/cc. CPF was purchased from Sigma-Aldrich as powder (purity > 98%). High-purity methanol (99.89%) was purchased from VWR. All the chemicals used for the determination of AChE activity were of the highest purity available.

#### 2.1.2 Model organism

Three-spined sticklebacks (*Gasterosteus aculeatus*) were collected in a reference site (Skaftö, Sweden [58°13’55.9“N 11°28’18.2“E]; water salinity: 16 – 18 ‰) with a hand-operated net, and immediately brought to the laboratory in aerated, thermally isolated boxes containing water from the sampling site. They were then acclimatized to and kept in artificial sea water for a month (13±1°C, 30 ‰, pH= 7.9) prior to start of the experiment. Fish were fed daily with red mosquito larvae. Continuous water flow and aeration ensured good water quality, and environmental enrichment was provided (gravel substrate pictures glued on the outer side of the tanks bottom). Fish were sexually mature, with a size range of 3.3 – 5.6 cm (median: 4.2 cm; average: 4.2 ± 0.5 cm; N = 96 individuals).

### 2.2 Spiking of microplastics with chlorpyrifos

CPF solubility in water is very low, therefore CPF dissolution was performed in methanol. To spike MPs, a CPF solution was prepared in methanol at the concentration of 30 mg/ml. Glass material was used for the spiking to limit chemical sorption on the walls of the vials. CPF solution was added to MP batches of approx. 100 mg, to obtain a final ratio between MPs and solvent (methanol) of 1:1 (w:w) (Smedes and Booij, 2012). The use of methanol to spike PE MPs was previously validated in a pilot experiment: no melting, color loss or other alteration of the MPs was observed (unpublished). MPs were left in contact with CPF solution for 10 days, and gentle agitation was provided to ensure homogeneous spiking of MPs. To force the partitioning of CPF to the particles, milli-Q water was gradually added during the whole spiking process to eventually reach 90% of the final volume. After the last day of spiking, MPs were filtered using 10 μm nylon filters, rinsed with 50 ml of milli-Q water, and filtered again for recovery. Five filtration-recovery cycles were performed before storage (4°C).

Five MP batches were prepared in total, each batch being sufficient for one week of organism contamination. A new batch of spiked MPs was prepared before every week of organismal exposure and used immediately for the exposure. Therefore, spiked MPs were stored for a maximum duration of one week. Analysis of spiked MPs subsamples shows an average CPF concentration of 17.7 ± 8.4 μg/mg (46.6% of CPF sorption, on average; see section 2.4 for quantification method).

### 2.3 Experimental setup

The exposure was carried out in September, once stickleback’s breeding season is over. Fish were exposed to pristine MPs (further referred to as “MPs”), CPF-spiked MPs (“MP-CPF”) or CPF only (“CPF”) via prey ingestion during four weeks. Exposure was performed by feeding individualized fish with pre-exposed prey every second day, three times a week, to limit handling of the fish. The number of Artemia fed to the fish increased over time, and a total of 51 *Artemia*/fish was reached at the end of the exposure (2 *Artemia*/fish/contamination day on weeks 1-2; 3 on week 3 and 5 on weeks 4-5). Every contamination day, live adult brine shrimps (*Artemia spp.*; approx. 1 cm) were placed for 15 minutes in Eppendorf tubes (2 ml; 15 *Artemia* individuals per tube) belonging to one of the experimental conditions (*i.e.* control, MPs, MP-CPF or CPF). Eppendorf tubes were prepared as follows, before the addition of *Artemia*: artificial seawater (ASW) only (control condition), approx. 2 mg of MPs or MP-CPF in ASW followed by strong manual shaking (MP and MP-CPF conditions, respectively), or CPF solution prepared in ASW (100 mg/L, solvent <10% total volume; CPF condition). After exposure, *Artemia* individuals were rinsed twice in ASW and fed to the fish from the respective experimental conditions less than five minutes later. Extra individuals were also kept and stored at −20°C after every contamination day, for further CPF concentration analysis. Exposure method is presented in figure 1.

**Figure 1:**
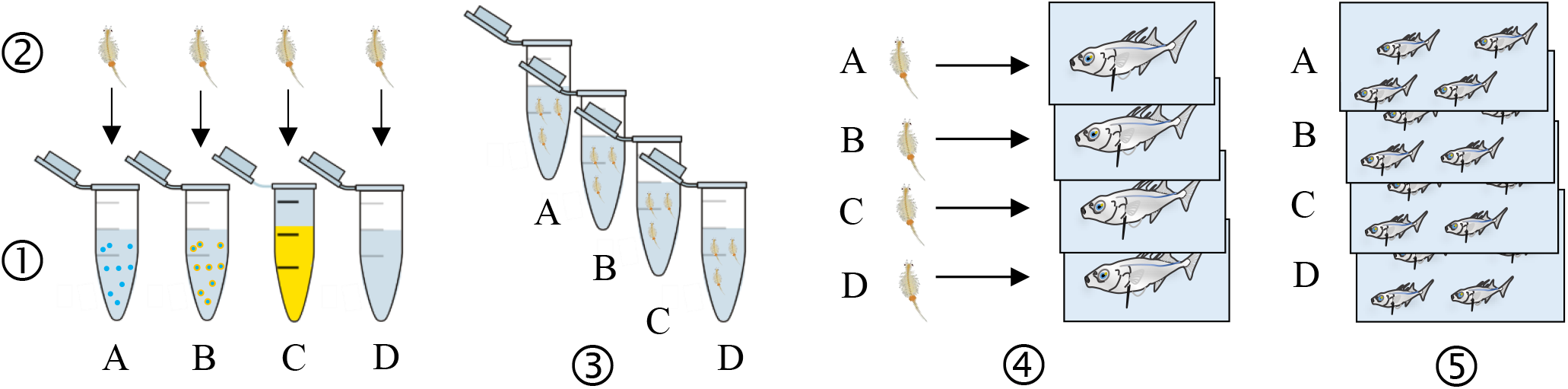
Schematic representation of the exposure method. Colors, organisms and MPs numbers and size are not accurate. 1) Eppendorf (n = 3) preparation for each condition: MPs in Artificial Sea Water (ASW) (A), MP-CPF in ASW (B), CPF in ASW (C) and control (ASW) (D). 2) Artemia (n = 15 per tube) are added and exposed for 15 min. 3) Exposure medium is removed and replaced by ASW to rinse Artemia (twice). 4) Artemia are fed to individualized fish (2 – 5 Artemia per fish. The number of Artemia increased over time, but was the same for every fish). 5) Fish belonging to the same exposure condition are grouped once all the Artemia have been ingested (8 fish per replicate, 3 replicates per condition).

Every experimental condition, including control, was performed in triplicates, each replicate (randomly allocated aquarium; 20L) comprising eight fish. Fish size range was 3.3 – 5.6 cm, with random allocation to exposure groups and no significant size differences between groups (ANOVA, p = 0.2). Exposure was performed under controlled temperature (14°C) and light (12:12, light:dark cycle) conditions. Half of the water was renewed every second day and aeration was provided in every aquarium, to ensure good water quality. Fish were individualized, during the feeding to ensure equal prey ingestion between fish, then placed back together in their respective aquaria. Outside exposure days, red mosquito larvae were provided *ad libitum* without individualization of the fish.

After two weeks of exposure, significant mortality was observed in fish from the CPF group. The exposure was therefore stopped for this group, behavior tests performed on both control and CPF groups, then fish from the CPF group were euthanized (48h after the last contamination). For MPs, MP-CPF and control groups, behavior assessment started after four weeks of exposure and lasted for a week. Exposure via prey continued during that week to prevent depuration of the fish and differing contamination levels between the first and last trial days. Behavioral trials were performed during the mornings and feeding in the afternoons, to prevent stress related to fish handling. After the last behavior trial (96h after the last contamination), fish were euthanized, measured, weighed and organs (gut, liver, gonads, muscle, gills and brain) were sampled for further analysis. Chemical analysis (CPF quantification) was performed on pooled fish that were found dead during the experiment (last contamination performed 48h before) and immediately stored at −20°C. Biochemical analyses (enzymatic activity, protein content) were performed on single, euthanized fish; samples were snap-frozen in liquid nitrogen and stored at −80°C.

### 2.4 Determination of microplastic ingestion and chlorpyrifos quantification

MPs selected for this study are blue beads that could easily be seen through *Artemia* cuticle under microscopic condition. Therefore, *Artemia* individuals (*n* = 47) were observed and photographed (Leica EZ4HD stereomicroscope with integrated HD camera) without prior sample preparation, immediately after their exposure and before being fed to the fish. Images were further analyzed with ImageJ^®^ software to count particles. Fish intestines (*n* = 17) were digested overnight in 10% KOH at 50°C (Bour et al., 2018). Extracts were then filtered on 10μm nylon mesh, filters observed under the stereomicroscope and particles counted.

CPF was quantified in MPs and organism samples by UHPLC-MS, using CPF-(diethyl-d10) as internal standard. Extraction and instrumental analysis were performed at the Swedish Environmental Research Institute (IVL). Additional information concerning chemical analysis can be found in supporting information.

### 2.5 Toxicity assessment

#### 2.5.1 Acetylcholinesterase (AChE) activity

AChE activity was determined in fish liver and brain, following an adapted procedure of Ellman’s method (Ellman et al., 1961; Sturve et al., 2016). Protein content was determined according to Lowry’s method (Lowry et al., 1951).

#### 2.5.2 Behavior assessment

Feeding, locomotion, environment exploration and reaction to the introduction of a novel object were assessed. For the feeding trial, fish were individualized, allowed to acclimatize for 10 minutes then fed two frozen mosquito larvae. The whole trial was video recorded and recordings were visually analyzed to determine the time required for each fish to ingest both larvae.

Locomotion, environment exploration and reaction to a novel object were assessed during a second trial, following the procedure described by Thompson et al. (Thompson et al., 2016). Fish were individualized in containers comprising one shelter (piece of tile) each, and allowed to acclimatize for 10 minutes. Video recording started after the acclimation period. After ten minutes of recording, a novel object (bolt attached to a transparent fishing line) was gently introduced in the center of the container and the fish reaction was recorded for 10 more minutes. Locomotion (immobility, total distance travelled, average speed, average acceleration, maximum speed, maximum acceleration) were determined over the first ten minutes of recording, using idTracker software. Time spent in shelter and fish behavior following the introduction of a novel object were determined by visual analysis. Fish containers were virtually divided between the area close to the shelter (half container) and the distal part of the container, to determine the total time spent inside the shelter, close to the shelter and far from the shelter, during the first 10 minutes. After introduction of the novel object, the assessed endpoints were (i) fish immediate reaction (*i.e.* freezing: immediate immobility and slight curving of the tail; escape: fast swimming on the opposite direction of the novel object; no specific reaction), (ii) delay in returning to normal behavior after the immediate reaction, (iii) delay in active observation of the novel object (*i.e.* fish facing the novel object), (iv) delay in approaching the novel object after active observation and (v) whether the fish actively touches the novel object or not.

### 2.6 Statistical analysis

All statistical analyses were performed using GraphPad Prism 8.1.1 software. Group comparisons of qualitative data (*i.e.* immediate reaction to a novel object and touching it or not) were performed with Chi-square tests. For quantitative data, normal distribution and homoscedasticity of residuals were verified with Shapiro-Wilk’s and Bartlett’s tests, respectively. Mann-Whitney tests were performed to compare CPF group and control group assessed after two weeks of exposure. One-way ANOVA tests were performed to compare MPs, MP-CPF and control group assessed after four weeks of exposure, when both normal distribution and homoscedasticity were verified. In cases of non-normal distribution, Kruskal-Wallis tests on rank were performed instead. Dunn’s test was performed to compare groups when significant differences were detected. Detailed data on the performed statistical analyses is presented in Supplementary Material (Supplementary Table 1). Levels of significance were set at p < 0.05. For behavior endpoints, trends were considered from p < 0.1.

## 3. Results

### 3.1 Trophic transfer of MPs and CPF

#### 3.1.1 MP ingestion and CPF accumulation in Artemia

*Artemia* ingested 204 ± 13 MPs/individual (average ± SE) before being fed to the fish. Based on MP ingestion and CPF concentrations in different MP batches, the expected (theoretical) CPF concentration in *Artemia* is 62.1 ng/individual. CPF quantification in *Artemia* samples from the MP-CPF group shows measured concentration of 97.3 ng/mg, equivalent to 57.8 ng/individual. Measured concentration in *Artemia* from the CPF group is 405.5 ng/mg, equivalent to 293.2 ng/individual.

#### 3.1.2 Trophic transfer of MPs and CPF in fish

Each fish was fed 51 *Artemia* in total, therefore ingesting theoretically 10 404 MPs in total (equivalent to 140 μg of MP). However, after four weeks of exposure MPs were found in only two fish (samples contained two and three particles, respectively).

CPF concentrations were measured in gonads, viscera (*i.e.* intestine, liver and gall bladder), body muscle, gills and brain (table 1). Daily checks ensured that dead fish were removed from the water and stored in less than 18 hours.

**Table 1:** CPF concentrations measured in fish organs from different exposure groups.

|  | CPF (ng/mg) |  |  |  |
| --- | --- | --- | --- | --- |
|  | Ctrl | MPs | MP-CPF<sup>1</sup> | CPF<sup>2</sup> |
| **Gonads** | < LOD | < LOD | 0.21 | 3.51 ± 0.21 |
| **Viscera** | < LOD | < LOD | 0.51 | 1.05 ± 0.20 |
| **Muscle** | < LOD | < LOD | 0.14 | 0.67 ± 0.06 |
| **Gills** | < LOD | < LOD | 0.07 | 0.49 ± 0.10 |
| **Brain** | < LOD | < LOD | 0.27 | 1.66 ± 0.09 |

<sup>1</sup>Values from one pool of fish (n = 3). <sup>2</sup>Average values (± SE) of two pools of fish (n = 4).

CPF transfer to fish via *Artemia* ingestion, both with and without the inclusion of MPs, was quite low (figure 2), with total values (addition of organ values) of 3% and 5.3%, for the MP-CPF and CPF groups, respectively. Single organ values ranged from 0.2 to 2% of total CPF ingested by the fish, depending on exposure conditions and organs. The relative distribution of CPF in the internal organs differed between MP-CPF and CPF groups (figure 3). While the organs showing the highest concentrations were the viscera and gonads for the CPF group (33 and 34%, respectively), most CPF was detected in the viscera (68%) in fish from the MP-CPF group. Compared to the CPF group, gonad samples from the MP-CPF group showed low percentage of CPF uptake (8%). Percentages indicated here correspond to the ratio (x100) between the total amount of CPF measured in organs (ng) and the total amount of CPF ingested via *Artemia* (ng; theoretical values).

**Figure 2:**
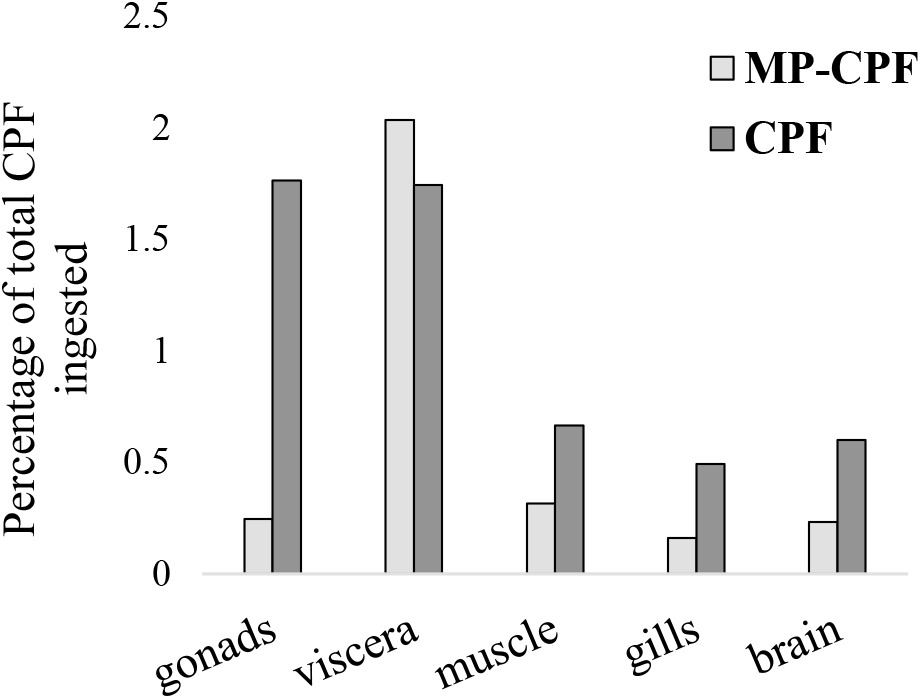
CPF transfer to fish, expressed as the ratio (x100) between the total amount of CPF measured in organs (ng) and the total amount of CPF ingested via *Artemia* (ng; theoretical values). CPF was quantified in one fish pool (n = 3) and two fish pools (n = 4), for the MP-CPF and CPF groups, respectively.

**Figure 3:**
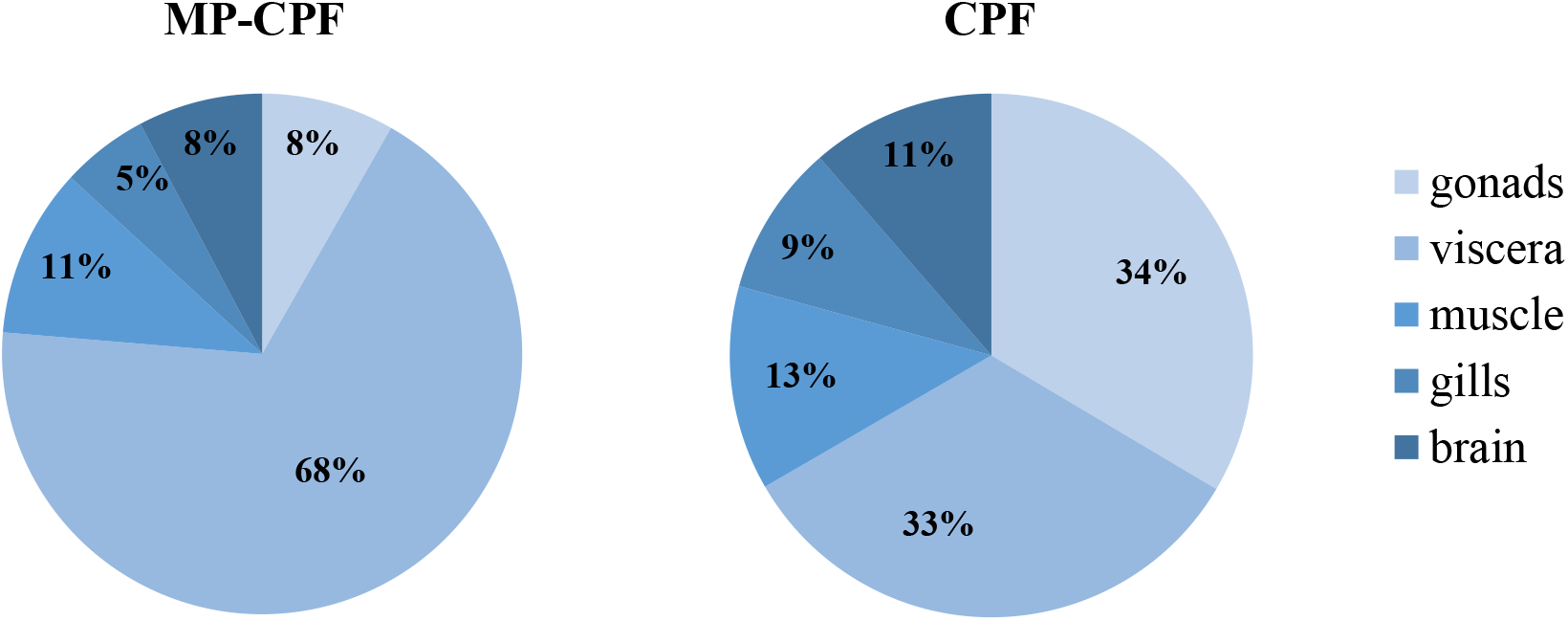
CPF relative distribution in different fish organs, expressed as the percentage of total CPF in fish. CPF was quantified in one fish pool (n = 3) and two fish pools (n = 4), for the MP-CPF and CPF groups, respectively.

### 3.2 Effects on fish

#### 3.2.1 Mortality

Significant mortality (58%) was observed in fish from the CPF group after two weeks of exposure. The exposure was therefore stopped for this condition after 16 days. The mortality recorded over the four weeks of exposure for control, MPs and MP-CPF groups was attributed to natural sensitivity of the species and handling stress, and considered non-significant (average mortality ± SE: 1.7 ± 0.3, 2.3 ± 0.9 and 2.7 ± 0.3 individuals, respectively).

#### 3.2.2 AChE activity

Decreased brain AChE activity was observed in fish exposed to MPs, MP-CPF and CPF (30.6%, 46.9% and 85.5% lower than control, respectively). However, the difference compared to the control group was statistically significant only for fish exposed to CPF alone (p < 0.01). In liver, 10% and 55% decreases were observed for the MP-CPF and CPF groups, respectively. An increase in liver AChE activity was observed in fish exposed to MPs (28.2% increase, compared to control). However, the differences observed between groups were not statistically significant (p > 0.05). AChE activities are presented in figure 4.

**Figure 4:**
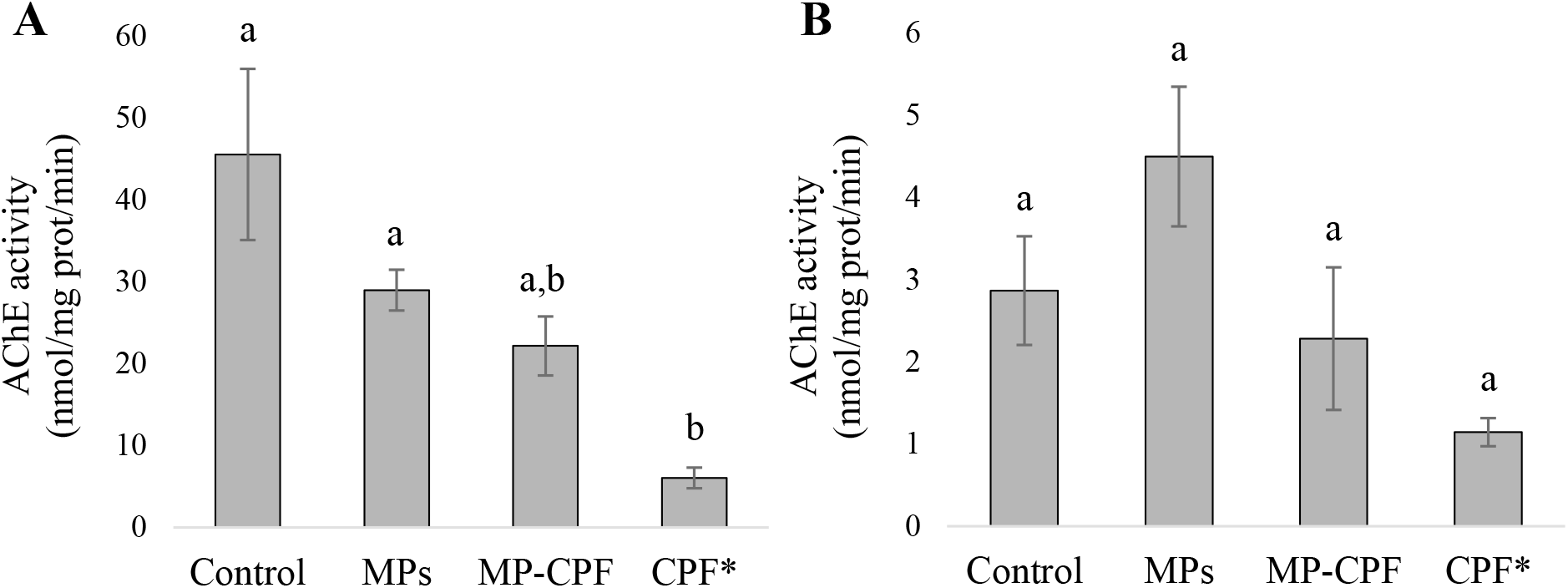
AChE activity (nmol/mg protein/min, mean values ± SE) in fish A) brain and B) liver. Different letters (a,b) indicate statistically different groups (p < 0.01). *Fish from the CPF group were euthanized and sampled after two weeks of exposure.

#### 3.2.3 Behavior

Behavior endpoints were grouped under four categories: feeding, locomotion, time spent in shelter and reaction to a novel object. Fish exposed to MPs did not present any behavioral changes compared to control fish. Fish exposed to MP-CPF exhibited changes in environment exploration and in their reaction to the introduction of a novel object. They spent less time in the shelter, compared to control fish (Δt = − 54 %; p = 0.1), and returned faster to a normal behavior after their first reaction following the introduction of the novel object (Δt = − 47 %; p = 0.1). Stronger behavioral changes were observed with fish exposed to CPF via prey, with changes in all four endpoint categories. CPF fish exhibited significantly longer feeding time and immobility (Δt = + 309 % and + 97%, respectively; p = 0.05), they spent less time in the shelter (Δt = − 73 %; p = 0.05) and active looking at and proximity with the novel object were delayed (Δt = + 138 % and 205 %, respectively; p = 0.05). Detailed results are presented in supplementary material (Supplementary Table S2, Supplementary Figures 1 and 2).

## 4. Discussion

### 4.1 Experimental trophic chain

Trophic transfer of MPs has been identified as a most relevant contamination pathway for predators (Lusher, 2015). In the present study, the selected trophic chain allows to control MP ingestion by preys and therefore fish exposure to MPs. We observed a consistent number of MPs in *Artemia* individuals throughout the experiment (204 ± 13 MPs per individual on average) and CPF concentrations measured in *Artemia* from the MP-CPF group (57.8 ng/individual) were very close to the expected concentrations (62.1 ng/individual). These results indicate that fish exposure to MPs and CPF was consistent throughout the experiment and validate the use of the present trophic chain as an appropriate method for controlled fish exposures. The absence of MPs in most fish samples is explained by the short particle retention time in three-spined stickleback (< 48h; Bour et al. 2020): the last contamination before fish were sampled was performed more than 48h and total egestion of MPs could be expected.

In the present study, the MP concentration used to expose *Artemia* (~1 mg/ml) is very high and not environmentally relevant. This concentration is not intended to represent realistic contamination conditions, but was chosen to maximize interactions between MPs and *Artemia.* Similarly, the average numbers of MPs ingested by *Artemia* and fish are too high to be environmentally relevant. A lower vector effect can therefore be expected in the environment. The MPs used for this study, pristine microspheres, were selected as model particles and do not represent the majority of MPs present in aquatic environments. The selection of these model particles can influence the observed effects: particle shape, specific surface area and the presence of other contaminants (*i.e.*”non-pristine” particles) highly influence sorption and desorption of chemicals (Heinrich et al., 2020), and therefore MP vector effect.

Chlorpyrifos is widely used in agriculture and in urban areas, and this compound and/or its metabolites are present in waters and sediments of streams, rivers, ponds, lakes and estuaries (Ware, 1999). The high concentration used to spike MPs is not representative of environmental concentrations (F. Müller et al., 2000; Marino and Ronco, 2005; Arain et al., 2018) but was chosen to compensate for potential loss of chemical during the spiking process (Smedes and Booij, 2012). For the same reasons, a high CPF concentration was used to contaminate *Artemia* (CPF group), as they were exposed in CPF solution for only 15 minutes. Pre-test were performed to ensure that these exposure conditions did not alter *Artemia* survival and swimming behavior (data not shown). Altered swimming ability of *Artemia* could have indeed influenced fish predation and resulted in fewer prey ingested.

### 4.2 Chlorpyrifos uptake in fish and vector effect of microplastics

CPF uptake varies markedly between organs (table 1), with bioaccumulation factors ranging from 0.001 (gills) to 0.009 (gonads) for fish from the CPF group. A previous study showed much higher bioaccumulation of CPF in *Aphanius iberus* exposed via contaminated *Artemia*, with a bioaccumulation factor of 0.3 (Varó et al., 2002). These results are not contradictory since CPF accumulation is highly dependent on fish species: other studies have investigated CPF accumulation in fish exposed via water and observed values ranging from 0.004 to 380 ng/mg (Thomas and Mansingh, 2002; Tilak et al., 2004; Rao et al., 2005). The detection of CPF in fish from the MP-CPF group shows that MPs can act as a vector for organic contaminants when ingested via the trophic chain. As ingested CPF quantities were initially different between MP-CPF and CPF groups, CPF transfer was expressed as the percentage of ingested CPF (figure 2). These values show that except in viscera, CPF accumulation in fish was much lower in the MP-CPF group. This result shows that although MPs can act as vector of contamination, contaminant transfer is limited compared to other exposure routes. The same phenomenon has been observed in most studies comparing contaminant uptake (PCBs, BFRs, PFCs, PBDEs, PAHs and organic contaminants) via MPs and other matrices (Browne et al., 2013; Grigorakis and Drouillard, 2018; Rainieri et al., 2018), although one study showed higher contaminant transfer in fish when exposed via MPs, compared to spiked food (Granby et al., 2018). Our results therefore confirm previous exposure and model studies, which concluded that MP vector effect could be negligible compared to natural pathways. This phenomenon has been explained by lower fugacity gradients between plastics and biota, compared to gradients between biota lipids (Koelmans et al., 2016). In our study, another factor could also contribute to this phenomenon: the digestion of natural prey likely resulted in the total release of CPF accumulated in prey tissue, while MPs were not digested, therefore limiting CPF release.

CPF accumulation varies between organs in both exposure conditions (figure 2). Previous studies showed CPF distribution patterns in fish similar to what was observed here in fish from the CPF group, gonads, brain and viscera being the most exposed organs (Thomas and Mansingh, 2002; Tilak et al., 2004). In fish exposed via MPs, CPF desorbed from the polymer and was mostly detected in the viscera. Batel et al. came to similar conclusions after exposing zebrafish via the trophic route to benzo[a]pyrene (BaP) sorbed on MPs: partial desorption of BaP from MPs was observed, with most of the BaP being detected in the intestinal tract and some detected in the liver, to a lower extent (Batel et al., 2016). Interestingly, the strongest BaP signal was detected in fish fed with Artemia contaminated via water, in line with a limited vector effect of MPs.

In other organs, initial sorption of CPF on MPs not only decreases CPF uptake, but also changes CPF distribution among organs (figure 3). While gonads seem to be the most exposed organ in the CPF group, the relative CPF concentration in gonads decreases in the MP-CPF group and reaches values below viscera and body muscle. This phenomenon can be explained by two different CPF release scenarios, either fast and total (CPF group) or low and constant (MP-CPF group), based on the combination of three factors: the low fugacity of CPF sorbed on MPs, the fast natural prey digestion versus the absence of digestion of MPs, and the degradation of CPF over time. Fish from CPF group rapidly digested their prey, which resulted in a fast and total release of CPF. This high CPF gradient between intestine and secondary organs resulted in fast and important transfer to secondary organs, especially gonads that are fat tissues and therefore accumulate hydrophobic contaminants more than muscle or gills. On the contrary, the low fugacity gradient of CPF sorbed on MPs resulted in low release. It can therefore be hypothesized that most CPF was degraded before reaching secondary organs. However, since MPs were not digested by fish, CPF release was constant and resulted in constant exposure of intestine, explaining the high concentrations found in this organ, in the MP-CPF group. CPF can be quickly degraded by organisms: in a previous study, total elimination of CPF was observed after only one day in fish exposed via the trophic route (Varó et al., 2002). Overall, our results suggest that when chemical exposure occurs via MPs, the organs most at risk can be different compared to exposure via water or via natural prey, decreasing gonads exposure in the case of CPF and dramatically increasing intestine exposure. However, fugacity gradients and release of sorbed chemicals depend on the properties of the considered chemical(s) (partition coefficient) and polymer(s) (binding capacity), and our findings might not hold true for every chemical-polymer combination. Studies involving different combinations of chemical and MP properties (partition coefficient and polymer type) are therefore needed to better understand MP vector effects.

### 4.3 Effect of contaminants

Similarly to our results, studies assessing the ecotoxicity of pristine PE MPs on fish exposed via trophic chains also reported no effects (Rochman et al., 2014; Mazurais et al., 2015; Jovanović et al., 2018). Moreover, most studies exposing fish via water showed either no effects (Ferreira et al., 2016; Karami et al., 2017; Batel et al., 2018; Malinich et al., 2018; Rainieri et al., 2018) or mild effects, which include inhibition of AChE activity, impaired energy reserves, decreased swimming and predatory performances, changes in plasma biochemistry and histological changes in gills (Oliveira et al., 2013; Luís et al., 2015; Karami et al., 2016; Choi et al., 2018; Wen et al., 2018). The adverse effects observed with pristine PE MPs in these studies could be due to the presence of monomers or additives (Lambert and Wagner, 2018) sorbed on MPs but that were not reported, either because no chemical analysis was performed or because concentrations were below limits of detection. Different hypotheses could explain the absence of effect following exposure via trophic chain, reported both in the scientific literature and in the present study. First, it could be explained by a faster elimination of MPs, directly related to gut retention time, while MPs could have a longer retention time when directly ingested or could get stuck in the gills when present in water. Another explanation could be that in trophic chain experiments, the additives potentially present on pristine MPs affect the preys but not the fish; the prey would therefore “protect” the predator species against the effects of MPs. One hypothesis is that the prey partially metabolizes the contaminant, thereby limiting the toxicity for predators.

In contrast with fish from the pristine MPs group, strong effects were observed on fish from the CPF group. As most organophosphate pesticides, CFP is a neurotoxic that inhibits AChE activity. Fish metabolize CPF to multiple metabolites, including CPF-oxon, which is the most efficient AChE inhibitor among the activated forms of CPF (Ware, 1999). It has been shown to be acutely toxic to fish, with 96h LC50 ranging from 300 to 650 μg/L (Tilak et al., 2004). The significant mortality observed here shows that CPF trophic transfer was important enough (up to 3.5 ng/mg; table 1) to induce acute toxicity in stickleback. In a previous study, authors reported no acute toxicity in Tilapia despite CPF accumulation values exceeding 100 ng/g (Thomas and Mansingh, 2002), which highlights important differences in species sensitivity to CPF. Unsurprisingly, significant brain AChE inhibition (84% decrease in activity) was observed in fish from the CPF group (figure 4). This strong inhibition is likely to be a cause of the important behavior impairment observed (Peakall et al., 2002). Taken together, the behavior changes observed show a dramatic hypoactivity, in comparison to control fish. Slow feeding is a sign of decreased predatory performance that could result in population changes for fish and their major prey (Weis et al., 2001). Increased immobility and time spent in the open field (the shelter being considered as a safe location) increase fish susceptibility to predation (Lafferty and Morris, 1996; Seppälä et al., 2004). Finally, fish behavior following the introduction of a novel object is indicative of their interaction with their environment. Previous studies observed changes in inspection of a novel object by fish, including stickleback, following exposure to chemicals (Maximino et al., 2010; Jutfelt et al., 2013). Here, the decreased curiosity, highlighted by the delay in observing and approaching the novel object, is likely to be a result of the severe hypoactivity in fish from the CPF group. These results show that CPF can severely impact stickleback behavior, with potential consequences at the population level. The reported behavior results should be interpreted with caution, given the small number of individuals left in the CPF condition.

While strong effects were observed in the CPF group with a general decrease in movement patterns, suggesting hypoactivity, fish exposed to CPF via MPs (MP-CPF) exhibited altered behavior to a lower extent, and in the opposite direction. Indeed, their faster reaction following the introduction of a novel object, combined with decreased time spent in the shelter, suggests hyperactivity. CPF has previously been shown to induce hyper excitability in exposed fish, increasing their vulnerability to predation (Little, 2002; Tierney et al., 2007; Halappa and David, 2009). It has also been shown that different behavior alterations occur at different thresholds of AChE inhibition (Tierney et al., 2007), resulting from different chemical exposure concentrations. This explains the differences in behavioral responses between CPF and MP-CPF exposed fish, since different levels of AChE inhibition were observed between the two groups (figure 4). These results show that the quantities of CPF transferred from MPs to fish are high enough to induce behavior impairment, potentially resulting in increased exposure to predation and increased energy expenditure, which should be considered when evaluating consequences at the ecosystem level (Galloway et al., 2017).

## Conclusions

Taken together, our results show that while PE MPs do not seem to cause adverse effects when ingested via prey, they can act as a vector for chemicals. Although lower transfer occurred when CPF was sorbed on MPs, compared to CPF accumulated in prey, it was important enough to induce adverse effects in stickleback. Moreover, CPF organ distribution in fish differed between exposure conditions, due to different CPF release scenario: either fast and total released from prey, or low and constant when released from MPs. Our results suggest that chemical exposure via MPs could alter the organ distribution of chemicals and result in a change in the organs most at risk, with a likely increase of intestine exposure.

## Ethics statement

Animal husbandry and experiments were conducted in compliance with ethical practices from the Swedish Board of Agriculture (Ethical permit number # 15986-2018).

## Authors contributions

AB, JS and BCA conceived the study; AB conducted the experiments; JH contributed to data analysis of fish behavior; AB wrote the manuscript, which all authors approved before submission.

## Acknowledgements

The authors are grateful to Dr. Georgios Giovanoulis (IVL) for his help with chemical characterization. We also acknowledge Dr. Tobias Lammel for his help with fish collection, and Magnus Lovén Wallerius for his help with setting up the video recording.

This manuscript has been released as a pre-print at bioRxiv, (Bour et al., 2019).

