## Supporting information_Bour et al for "Microplastic vector effects: are fish at risk when exposed via the trophic chain?" (from another version of this preprint: version 1, 2019-12-18)

Total number of pages: 9

Total number of tables: 2

Total number of figures: 2

### Table of contents

**Supplementary methods: Chlorpyrifos quantification** **S3**

**Table S1.** Statistical tests performed and p-value results. **S4**

**Table S2.** Behavior assays results. **S7**

**Figure S1.** Distribution of behavioral responses of fish exposed to microplastics (MPs) and spiked MPs (MP-CPF). **S8**

**Figure S2.** Distribution of behavioral responses of fish exposed to chlorpyrifos (CPF). **S9**

### Supplementary methods: Chlorpyrifos quantification

#### *Extraction Method*

Each sample was spiked with 100 ng Chlorpyrifos-(diethyl-d10), CAS number 285138-81-0, as internal standard (IS) obtained from Sigma-Aldrich. Samples were extracted with ultra-sonication for 15 min and centrifugation at 4000×g for 5 min using 2 x 2 mL MeOH and 2 x 2 mL ACN. The four supernatants from each extraction were combined and each sample was evaporated to a final volume of 0.5 mL ACN. Extracts were left at -20°C until the analysis.

#### *Instrumental analysis*

Chloropyrifos, CAS Number 2921-88-2 (Pestanal, analytical standard from Sigma-Aldrich) and chlorpyrifos-(diethyl-d10) were analyzed with UHPLC Dionex Ultimate 3000 (Thermo Fisher Scientific) coupled with a Thermo Scientific Q Exactive Focus Hybrid Quadrupole-Orbitrap Mass Spectrometer using heated electron spray ionization (+HESI) in positive ion polarity mode. Chromatographic separation was carried out on analytical dC18 Atlantis (2.1 × 150mm, 3.0µm particle size) column (Waters) equipped with a pre-column (VanGuard, Waters) using a nominal flow of 0.6 ml/min. The mobile phase A was 10 mM $CH_3COOH$ in $H_2O$ and mobile phase B 10 mM $CH_3COOH$ in MeOH. The total run time was 12 min and the gradient elution set as follows: 0-1 min 30% B, 1-8 min 95% B, 8-10.5 min 95% B, 10.5-11 min 30% B, 11-12 30% B. Parameters for the heated electrospray ionization (+HESI) were the following: sheat gas flow rate 48, aux gas flow rate 11, sweep gas flow rate 2, spray voltage 3.5kV, capillary temp 256°C, S-lens RF level 50, aux gas heater temp 413°C. Pierce LTQ Velos ESI positive ion calibration solution (ThermoFisher Scientific) was used for positive mode calibration of the orbitrap each time before the analysis.

The samples were analyzed in full scan - data depended $MS^2$ (FS-dd$MS^2$) using a scan range of 320-400 m/z. Full scan had resolution of 70 000 FWHM, and dd-$MS^2$ for confirmation had resolution of 17 500 FWHM, isolation window 3.0 m/z, N(CE) 10. 30, 50, maximum IT 50 msec and apex trigger 3 to 8 sec. The inclusion list for the data depended dd-$MS^2$ confirmation in positive polarity included the masses (m/z) 349.9335 and 351.9306 for chlorpyrifos, and 359.9963 and 361.9934 for chlorpyrifos-(diethyl-d10) internal standard.

All samples were quantified using the Thermo TraceFinder 4.1 software for Environmental and Food Safety (EFS).

**Table S1:** Statistical tests performed and p-value results.

| Endpoint | Tested groups | Test performed | p-value | Post-hoc test performed | Post-hoc test adjusted p-value |
|---|---|---|---|---|---|
| **Brain AChE** | Ctrl (n = 12)<br>MPs (n = 13)<br>MP-CPF (n= 9)<br>CPF (n = 8) | Kruskal-Wallis | p = 0.002 | Dunn's | Ctrl vs MPs: p > 0.99<br>Ctrl vs MP-CPF: p = 0.8<br>Ctrl vs CPF: p = 0.001<br>MPs vs CPF: p = 0.01<br>MPs vs MP-CPF: p > 0.99<br>MP-CPF vs CPF: p = 0.24 |
| **Liver AChE** | Ctrl (n = 9)<br>MPs (n = 11)<br>MP-CPF (n= 8)<br>CPF (n = 6) | Kruskal-Wallis | p = 0.44 | none | |
| **Feeding time** | Ctrl (n = 17)<br>MPs (n = 17)<br>MP-CPF (n= 18) | Kruskal-Wallis | p = 0.26 | none | |
| | Ctrl-CPF (n = 18)<br>CPF (n = 9) | Mann-Whitney | p = 0.003 | none | |
| **Immobility** | Ctrl (n = 19)<br>MPs (n = 18)<br>MP-CPF (n= 19) | Kruskal-Wallis | p = 0.44 | none | |
| | Ctrl-CPF (n = 12)<br>CPF (n = 6) | Mann-Whitney | p = 0.018 | none | |
| **Total distance** | Ctrl (n = 19)<br>MPs (n = 18)<br>MP-CPF (n= 19) | Welch' ANOVA | p = 0.43 | none | |
| | Ctrl-CPF (n = 12)<br>CPF (n = 6) | Mann-Whitney | p = 0.75 | none | |
| **Average speed** | Ctrl (n = 19)<br>MPs (n = 18)<br>MP-CPF (n= 19) | Kruskal-Wallis | p = 0.62 | none | |
| | Ctrl-CPF (n = 12)<br>CPF (n = 6) | t-test | p = 0.62 | none | |

| | | | | |
|---|---|---|---|---|
| **Average acceleration** | **Ctrl (n = 19)**<br>**MPs (n = 18)**<br>**MP-CPF (n= 19)** | **Kruskal-Wallis** | **p = 0.52** | **none** |
| | Ctrl-CPF (n = 12)<br>CPF (n = 6) | t-test | p = 0.75 | none |
| **Max. speed** | Ctrl (n = 19)<br>MPs (n = 18)<br>MP-CPF (n= 19) | Kruskal-Wallis | p = 0.20 | none |
| | Ctrl-CPF (n = 12)<br>CPF (n = 6) | t-test | p = 0.97 | none |
| **Max. acceleration** | Ctrl (n = 19)<br>MPs (n = 18)<br>MP-CPF (n= 19) | Kruskal-Wallis | p = 0.36 | none |
| | Ctrl-CPF (n = 12)<br>CPF (n = 6) | t-test | p = 0.94 | none |
| **Time spent in the shelter** | Ctrl (n = 19)<br>MPs (n = 18)<br>MP-CPF (n= 19) | Kruskal-Wallis | p = 0.10 | none |
| | Ctrl-CPF (n = 12)<br>CPF (n = 6) | t-test | p = 0.029 | none |
| **Time spent far from the shelter** | Ctrl (n = 19)<br>MPs (n = 18)<br>MP-CPF (n= 19) | ANOVA | p = 0.36 | none |
| | Ctrl-CPF (n = 12)<br>CPF (n = 6) | t-test | p = 0.029 | none |
| **Return to normal behavior** | Ctrl (n = 16)<br>MPs (n = 12)<br>MP-CPF (n= 12) | Kruskal-Wallis | p = 0.06 | none |
| | Ctrl-CPF (n = 10)<br>CPF (n = 7) | Mann-Whitney | p = 0.19 | none |
| **Delay in active observation of NO** | Ctrl (n = 19)<br>MPs (n = 17)<br>MP-CPF (n= 17) | Kruskal-Wallis | p = 0.23 | none |
| | Ctrl-CPF (n = 12)<br>CPF (n = 4) | Mann-Whitney | p = 0.018 | none |

| | | | | |
|---|---|---|---|---|
| **Delay in NO approach** | **Ctrl (n = 19)**<br>**MPs (n = 18)**<br>**MP-CPF (n= 15)** | **Kruskal-Wallis** | **p = 0.16** | **none** |
| | Ctrl-CPF (n = 11)<br>CPF (n = 4) | Mann-Whitney | p = 0.037 | none |

**Table S2:** Behavior assays results.

| | | Control 1 | Control 2 | MPs | MP-CPF | CPF |
|---|---|---|---|---|---|---|
| **Feeding** | feeding time (s) | 70 ± 15 | 42 ± 10 | 38 ± 14 | 56 ± 14 | 286 ± 80** |
| **Locomotion** | immobility (s) | 36 ± 07 | 50 ± 05 | 71 ± 17 | 70 ± 17 | 71 ± 11** |
| | total distance[1] | 107 ± 30 | 29 ± 01 | 28 ± 03 | 33 ± 03 | 96 ± 27 |
| | average speed[2] | 164 ± 43 | 32 ± 01 | 30 ± 02 | 32 ± 03 | 155 ± 28 |
| | average acceleration[3] | 35 ± 08 | 7 ± 0.3 | 6 ± 0.4 | 7 ± 0.6 | 33 ± 07 |
| | max. speed[2] | 6615 ± 662 | 711 ± 134 | 945 ± 499 | 627 ± 96 | 6674 ± 1449 |
| | max. acceleration[3] | 1586 ± 166 | 163 ± 34 | 227 ± 125 | 146 ± 25 | 1561 ± 363 |
| **Environment exploration** | time spent in the shelter (s) | 199 ± 41 | 94 ± 22 | 90 ± 35 | 43 ± 13* | 53 ± 24** |
| | time spent far from the shelter (s) | 223 ± 46 | 281 ± 35 | 331 ± 40 | 351 ± 33 | 230 ± 60 |
| **Reaction to a novel object (NO)** | immediate reaction | freeze (6)<br>escape (8)<br>no reaction (2) | freeze (9)<br>escape (7)<br>no reaction (3) | freeze (5)<br>escape (7)<br>no reaction (6) | freeze (7)<br>escape (5)<br>no reaction (7) | freeze (2)<br>escape (5)<br>no reaction (0) |
| | return to normal behavior (s) | 69 ± 26 | 51 ± 12 | 33 ± 10 | 27 ± 06* | 137 ± 73 |
| | delay in active observation of NO (s) | 124 ± 32 | 149 ± 05 | 128 ± 30 | 126 ± 26 | 355 ± 87** |
| | delay in NO approach (s) | 36 ± 20 | 41 ± 19 | 46 ± 23 | 33 ± 17 | 125 ± 39** |
| | NO touching | yes (4)<br>no (12) | yes (4)<br>no (15) | yes (6)<br>no (12) | yes (3)<br>no (16) | yes (1)<br>no (6) |

Quantitative data are presented as mean values ± standard error. Qualitative data are followed by the number of individuals associated to the corresponding data (numbers in brackets).

[1]expressed as "unit" (relative values)

[2]expressed as "unit/s" (relative values)

[3]expressed as "unit/$s^2$" (relative values)

"Control 1" and "Control 2" are results from the control group, assessed after two and four weeks of exposure, respectively.

Group comparison was performed as follow: (i) "CPF" compared to "Control 1" (Mann-Whitney test); (ii) "MPs" and "MP-CPF" compared to "Control 2" (ANOVA/Kruskal-Wallis tests).

*statistically different from "Control 2" ($p < 0.1$)

**statistically different from "Control 1" ($p < 0.05$)

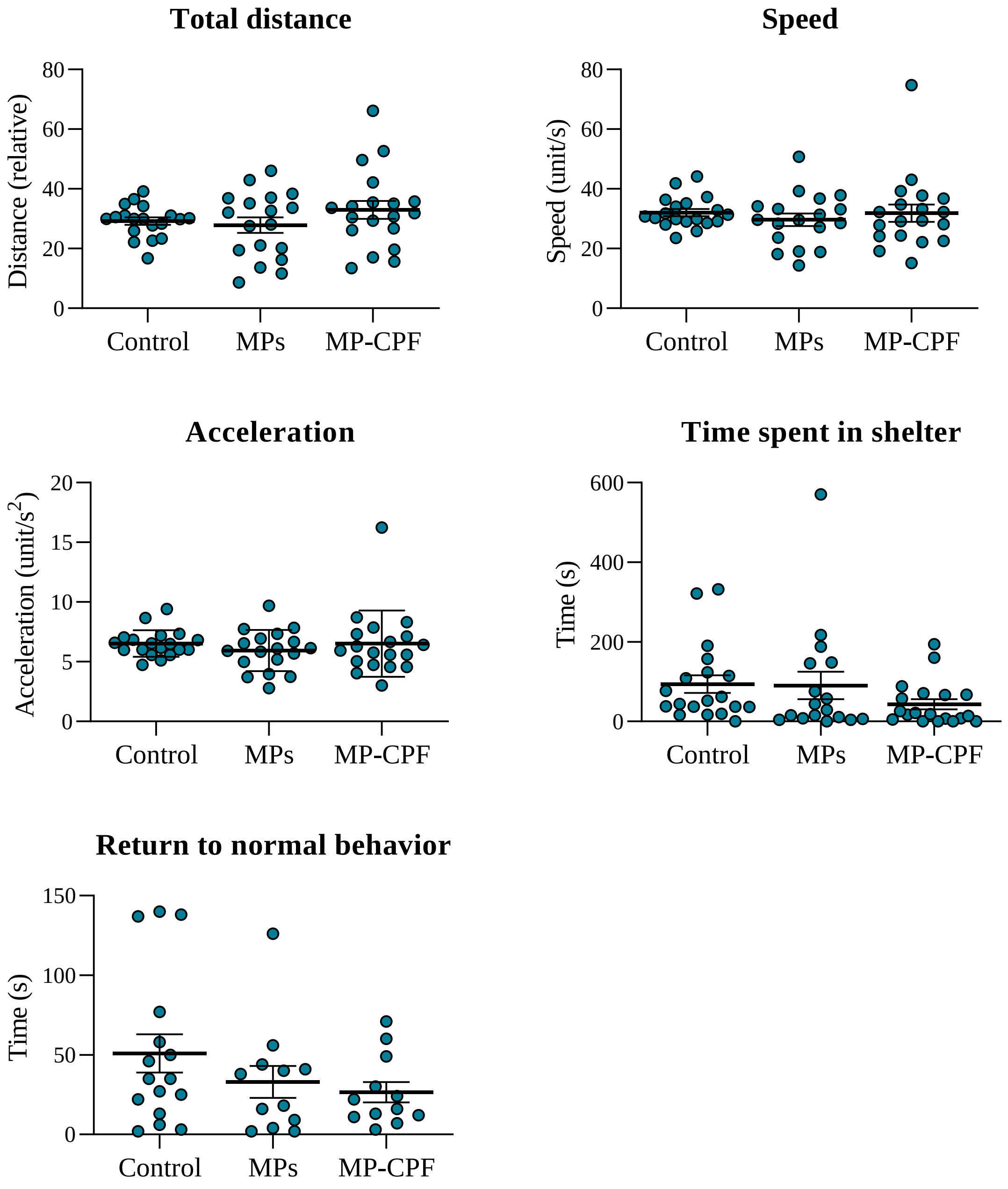


**Figure S1:** Distribution of behavioral responses of fish exposed to microplastics (MPs) and spiked MPs (MP-CPF). Each dot represent a single response (single fish). Mean values and standard errors are represented as thick and thin bars, respectively.

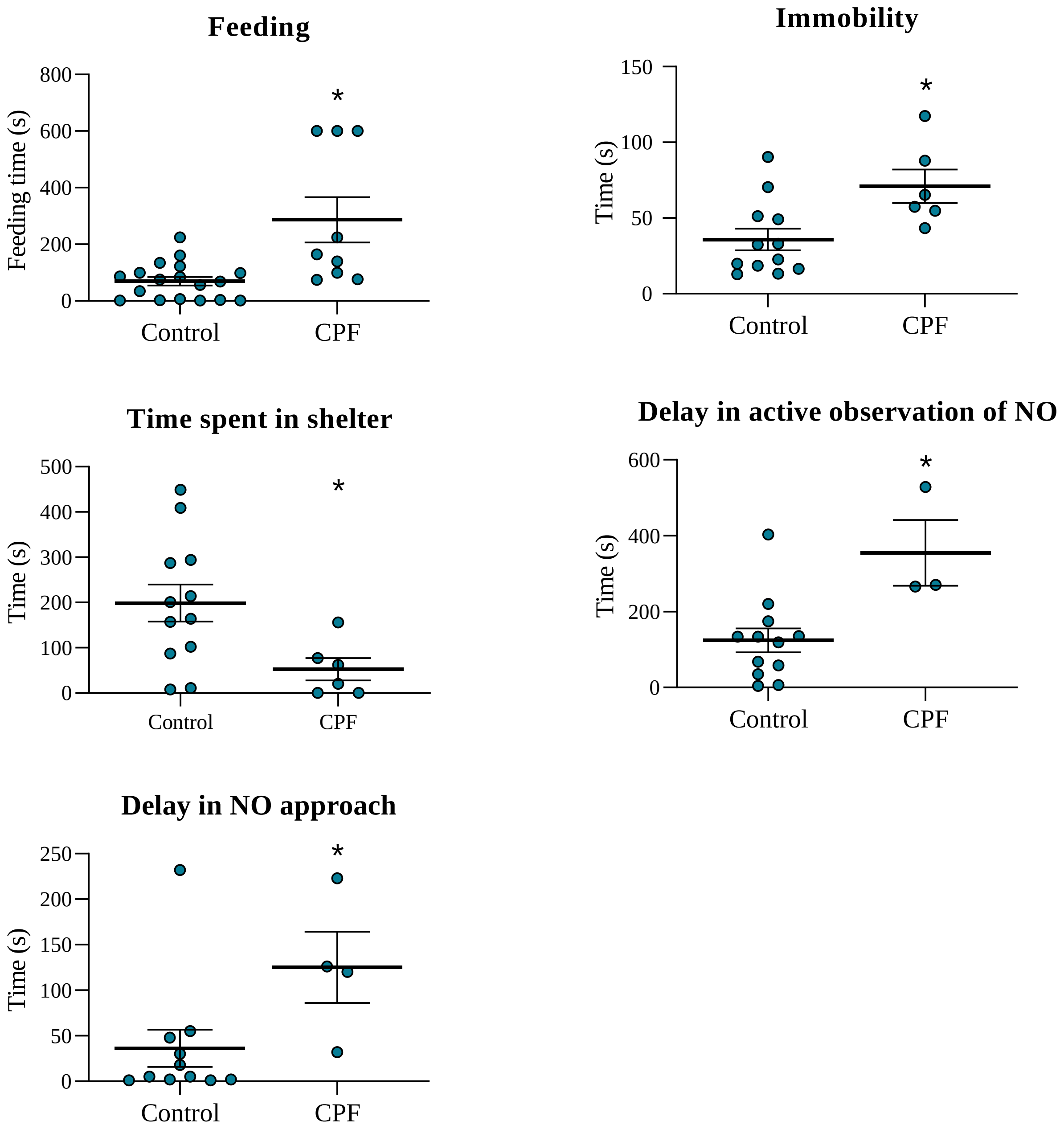


**Figure S2:** Distribution of behavioral responses of fish exposed to chlorpyrifos (CPF). Each dot represent a single response (single fish). Mean values and standard errors are represented as thick and thin bars, respectively. Selected endpoints present significant difference ($p < 0.05$) between CPF and Control groups, indicated by asterisks (*). NO: Novel Object.
